# Building RNA Backbone in Constant Time by Numerical Approximation

**DOI:** 10.1101/2023.10.17.562784

**Authors:** Philippe Thibault, François Major

## Abstract

We present a numerical approximation method designed to swiftly create RNA ribose conformations in constant time. The method’s parameterization relies on the atomic coordinates of a given base along with its two neighboring phosphate groups. Such a parameterization suits three dimensional modeling engines that determine the RNA conformational search space using base operations instead of backbone sampling. These engines consider phosphate groups as part of the base rigid bodies. Reconstructing ribose conformations result in less than 1 Å of RMSD (root-mean-square deviation) compared to original conformations derived from high-resolution X-ray crystallographic structures. By incorporating this ribose construction method into *MC-Sym*, a well-established RNA three dimensional modeling software, we streamline the modeling process into two phases. This enhances the search algorithm’s speed and improves model consistency and precision. Additionally, we employed the method to pinpoint 27 irregular ribose stereoisomers in high-resolution RNA X-ray crystal structures.

## Introduction

RNAs are made up of nucleotides. Each nucleotide consists of a nitrogenous base (either adenine, guanine, cytosine, or uracil), a sugar molecule (the ribose), and a phosphate group (1). In the RNA chain, phosphate groups and riboses alternate, with a nitrogenous base attached to each ribose. This sequence of phosphate and ribose forms what is commonly known as the backbone or phosphodiester linkage of the RNA. **Figure 1A** provides a three dimensional (3D) depiction of a tri-nucleotide RNA, emphasizing its bases, riboses, and phosphate groups. Geometrically speaking, the RNA backbone is notably flexible and is defined by eleven free torsion angles. Capturing this inherent flexibility accurately is a significant challenge when computationally modeling RNA in 3D. There have been numerous attempts to produce distinct sets of nucleotide conformations by clustering them based on coordinates (2–4) and torsion angles (5–10).

**Figure 1.**
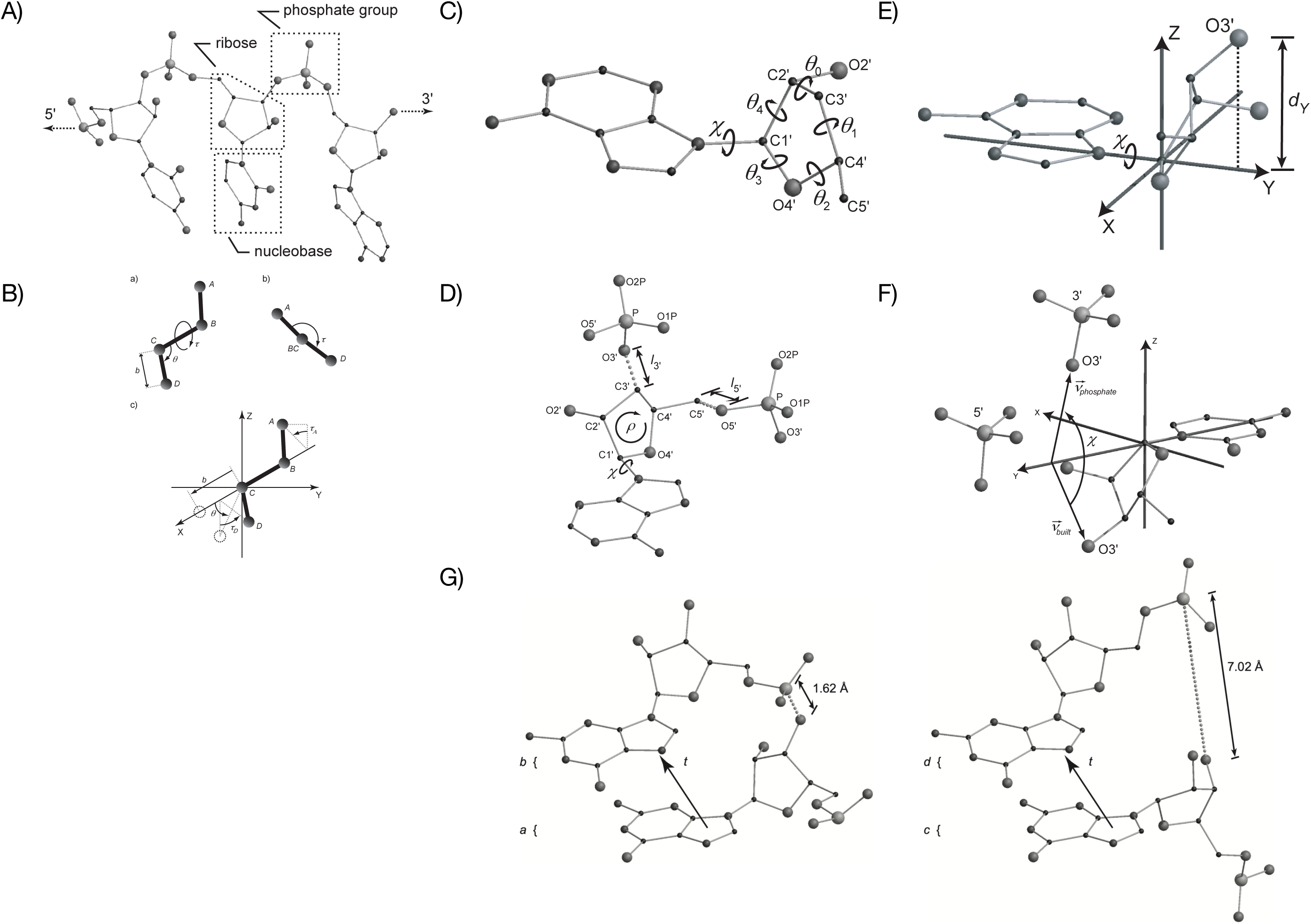
Structural components and properties of a nucleotide. **A**) Base, ribose and phosphate group. A strand of three nucleotides is shown, linking in order an uracil, a cytosine and an adenine with their respective riboses and phosphate groups. The dashed arrows indicate the linkage to the O5’ of the preceding nucleotide (left) and to the P of the succeeding one (right). The radius of the spheres is proportional to the mass of the represented atoms (C < N < O < P). The hydrogen atoms are not shown. **B**) Covalent bond geometry. a) Bond length, bond angle, and torsion angle. Given four atoms *A*, *B*, *C* and *D*, the bond length *b* between atoms *C* and *D* is the length of vector *CD*. The bond angle θ between bonds *BC* and *CD* is the angle between vectors *BC* and *CD*. The torsion angle τ between atoms *A* and *D*, around bond *BC*, is the angle between the projection of vectors *AB* and *CD* on a plane orthogonal to vector *BC*. b) Computation of torsion τ. *A’*, *B’*, *C’*, and *D’* are the projections of respectively *A*, *B*, *C* and *D* to a plane orthogonal to *XYZ* (paper plane). The torsion τ is the angle between *A’B’* and *C’D’*. c) Computation of atom D, given *A*, *B*, and *C*. The coordinate frame is aligned with bond *BC* and the origin at *C*. In this frame, the length of bond *CD* is expressed by a translation of magnitude *b* along the axis aligned with bond *BC* (X axis). Then, the bond angle between *BC* and *CD* is expressed by a rotation of angle θ around an axis orthogonal to bond *BC* (Y axis). Finally, the torsion angle between *A* and *D*, around *BC*, is expressed by a rotation of τD around the axis aligned with bond *BC* (X axis), where τD = 180o + τA – τ. The composition of the three above transformations moves *D* from the frame’s origin to its final location, expressing the covalent geometry defined in (a). **C**) Torsion angles and atomic numbering of the ribose. Here a ribose is shown bound to an adenine. **D**) Objective function. A base (here adenine) and two phosphate groups are shown with a ribose built in between from the input parameters ρ and χ. The return value is the average error in the length of bonds C5’O5’ and C3’O3’, respectively l5’ and l3’ shown with dotted lines. **E**) Measurement of *dY*: the distance between O3’ and the Y axis in a coordinate frame aligned onto the glycosyl bond. In this frame, χ is a rotation about the Y axis, thus *dY* is independent of χ. **F**) Estimation of χ. Given the positions of a base and two phosphate groups, Vbuilt is orthogonal to the glycosyl bond and passes through the O3’ atom from the 3’ phosphate. vphosphate is orthogonal to the glycosyl bond and passes through the O3’ atom that is positioned as an exocyclic substituent onto an intermediate ribose (χ = 0°). Then, the estimated value for χ is the angle between these two vectors. **G**) Two models of stacked G nucleotides. a) Selection of G nucleotide conformations *a* and *b* and of transformation matrix *t*, results in a model where the interconnecting O3’P bond is of acceptable length (1.6 Å). b) Using the same stacking transformation matrix as in (a), *t*, but two different nucleotide conformations – *c* and *d* – the O3’P bond is stretched to 7.0 Å. This model can be difficult to repair by numerical methods, even if the guanines are properly stacked.

If we operate under the assumption that RNA folding is primarily influenced by hydrogen-bonded base pairing and dipole-induced base stacking, it logically follows that the 3D configuration of the backbone is tailored to accommodate these base pairing and stacking patterns. In our latest advancements with the RNA 3D modeling software, *MC-Sym* (11,12), we first focused on a conformational search space determined by the spatial positioning of the bases. Only after that did we address the fitting of the backbone using numerical approximations. Within this framework, we devised a method that constructs the ribose structure based on the 3D coordinates of a specific base and its neighboring phosphate groups. In this system, each base, along with its associated phosphate group (13), is treated as a single rigid entity. The interactions due to base pairing and stacking are captured using rigid body transformations (14).

The task of constructing a ribose conformation that connects a base with two phosphate groups can be formally described as an optimization challenge. To address this, we have implemented two optimization techniques to precisely determine the ribose conformation: the derivative-less linear optimization and a constant-time estimation method. We evaluated the efficacy of these methods using a benchmark that involves reconstructing riboses from high-resolution X-ray crystallographic data. Our findings indicate that integrating a ribose construction method into MC-Sym considerably enhances its performance in terms of speed, precision, and consistency. Additionally, we employed our method to examine the stereochemistry of riboses present in high-resolution X-ray crystallography models.

## Methods

In this section, we detail the ribose construction method, framing it as an optimization problem. We start with the given spatial coordinates of a base and its neighboring two phosphate groups. To guide the construction, we implemented an objective function. This function constructs the ribose configuration and then measures the average discrepancy in the implicit covalent bonds connecting the ribose to both phosphate groups. This discrepancy is quantified using the root-mean-square deviations (RMSD) from the ideal values. We begin by formally defining this objective function and its parameters. Subsequently, we introduce two algorithms designed to optimize this function: one utilizes a derivative-less linear optimization technique, while the other employs constant time optimal parameter estimation.

### Objective function

The objective function’s purpose is to construct a ribose conformation based on its parameters, which represent the degrees of freedom of the ribose conformation. In our approach, we treated all covalent bonds and angles as having fixed optimal values, with only the torsion angles left as variables; these torsion angles form the parameters for the objective function. The ideal values for the covalent bonds and angles of a ribose structure were sourced from the NDB website (15). These values were determined through numerical optimization using the X-PLOR program (16).

Upon analyzing the structural configurations of nucleotides, it became evident that the covalent geometry of the pentose sugar aligns with discernible stereochemical conformations (17). This means that if we set the covalent geometry to a specific torsion angle value during the construction process, the resulting structure will inevitably exhibit some deformations. To measure these deviations, we curated a database, Σ. This database compiles RNA X-ray crystallographic structures with a resolution of 2.4 Å or finer, sourced from the RCSB’s PDB (18) (Research Collaboratory for Structural Bioinformatics Protein Data Bank) up to June 2005 (refer to **Supplementary Table** for the list of selected PDB entries). From this database, we calculated the average discrepancies between the bond lengths and bond angles of the 17,749 ribose configurations and their optimal values. With the obtained minimal mean deviations - 0.006 Å for bond length and 0.974° for bond angle – it became evident that maintaining a fixed covalent geometry for the ribose is a reasonable approximation. Thus, focusing on free torsion angles as the primary parameters proves adequate for ribose construction.

#### Torsion angles

Torsion angles provide insight into the spatial arrangement of four sequential atoms – A, B, C, and D. Specifically, they indicate how the bonds A-B and C-D orient relative to the central bond B-C (as visualized in **Figure 1B**ab. Given the positions of atoms A, B, and C, atom D’s location can be deduced by considering three factors: the length of the bond C-D, the angle formed between bonds B-C and C-D, and the torsion revolving around bond B-C. The optimal lengths and angles for bonds are determined by the standard covalent geometry. The exact placement of atom D hinges on the torsion surrounding bond B-C, which remains a variable. **Figure 1B**c graphically demonstrates how the positioning of atom D, in relation to atoms A, B, and C, establishes the free torsion around bond B-C.

Figure 1C illustrates the torsion angles that influence the conformation of the ribose: there is one associated with the glycosyl bond (χ) and five around the furanose ring, labeled as θ0 through θ4. Utilizing the aforementioned construction procedure, each atom in the ribose is positioned based on a specific torsion parameter.

To begin, the C1’ atom is anchored to the base using a predetermined covalent geometry, thereby forming the glycosyl bond. Proceeding from there, and using a frame of reference set by the glycosyl bond, the positions of atoms C2’ and O4’ are derived from C1’ based on the torsion angle χ. By shifting the frame along the bond between C1’ and C2’, atom C3’ is placed using the torsion parameter θ4. To complete the ring structure, atom C4’ is aligned in relation to the C1’-O4’ bond, guided by the torsion angle θ2. Finally, the external atoms C5’ and O2’ are connected to the already constructed ring, rounding off the full conformation of the ribose.

#### Parameters

In 1972, Altona and Sundaralingam streamlined the torsion space associated with a furanose ring. They reduced it from a 5-dimensional representation – denoted as <θ1, θ2, θ3, θ4, θ5> – to a simpler 2-dimensional form – symbolized as <ρ, θm>. Here, ρ represents the phase angle of the ring’s pseudorotation (originally labeled as P), while θm indicates its amplitude, which is the highest possible torsion value any torsion angle of the ring can achieve (19). This dimensional reduction stems from the interrelatedness of the furanose ring’s torsion angles, a consequence of its cyclic nature and the exocyclic components attached to it. The correlation between <θ1, θ2, θ3, θ4, θ5> and <ρ, θm> can be expressed through **Equation 1**, where *i* can take any value from the set {0, 1, 2, 3, 4} and Δ is a constant, equal to 144°.

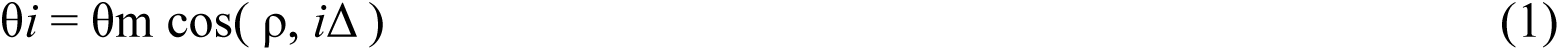

Equations 2 and 3 are derived from Equation 1 to obtain <ρ, θm> from <θ1, θ2, θ3, θ4, θ5>.

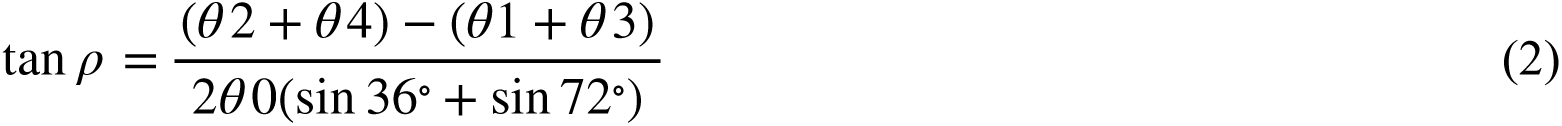

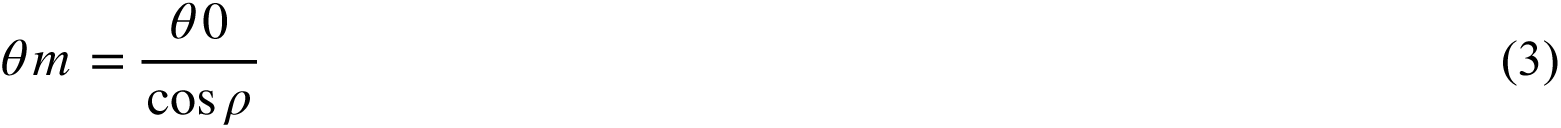

By applying Equation 1, the torsion angles <θ1, θ2, θ3, θ4, θ5> can be derived from <ρ, θm>. These angles are instrumental in constructing the furanose ring that is connected to a base via the torsion χ surrounding the glycosyl bond. In our approach, we set θm to its value as observed in the ideal ribose configuration, which is θm = 37.68°. This leaves both ρ and χ as the only adjustable parameters in our construction method.

It is worth noting that this construction technique serves as our objective function, aimed at optimization. This function can be expressed as outlined in Equation 4. Here, l5’ and l3’ respectively represent the lengths of the implicit bonds C5’-O5’ and C3’-O3’. Meanwhile, λ5’ and λ3’ are the corresponding lengths as seen in the ideal ribose, with values λ5’ = 1.440 Å and λ3’ = 1.431 Å (Figure 1D).

Consequently, pinpointing the best parameters to minimize the function *f*(ρ, χ) corresponds to forming a ribose conformation connected to a base. The goal is to achieve the smallest RMSD between the lengths of the implicit bonds connecting it to the phosphate groups and their optimal, pre-defined lengths.

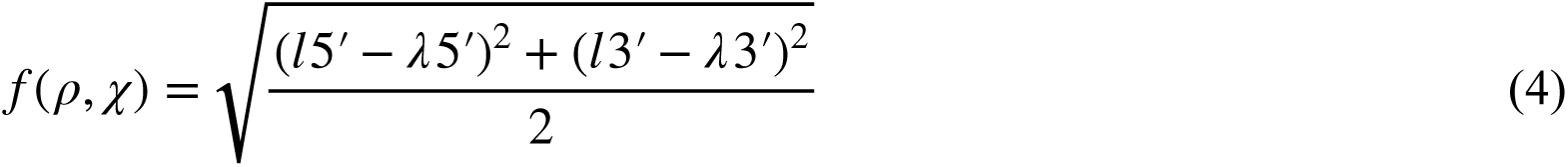

#### Precision

At this juncture, the precision of our construction technique is a primary concern. This is because the objective function, *f*, does not explicitly address certain geometrical attributes of a ribose. Notably, these include the C3’-C4’ bond, which completes the ring once its five atoms have been placed, and the torsions θ0, θ1, and θ2, which are not directly utilized in determining the positions of the ring’s atoms.

In **Figure 2A** (left), we examined the C3’-C4’ bond length for riboses built across a complete 360° ρ value range. We then juxtaposed the deviations of this length against the corresponding length observed in a reference structure. Our analysis revealed that, depending on the position within the pseudorotation cycle, the closing bond’s length could be shorter by as much as 0.049 Å or extend longer by up to 0.123 Å. Such variations represent the inherent distortions we encounter in ribose conformations using this method.

To gain a deeper understanding of these distortions, Figure 2A (right) visually maps the furanose’s five torsion angle values against the associated pseudorotation over a complete cycle. By comparing the solid and dotted lines in the figure, we can gauge the discrepancies between the torsion values derived from **Equation 1** and those from the constructed ribose. The close correspondence between the theoretical and actual values underscores the minimal errors attributable to deviations in the standard covalent geometry of furanosides.

**Figure 2.**
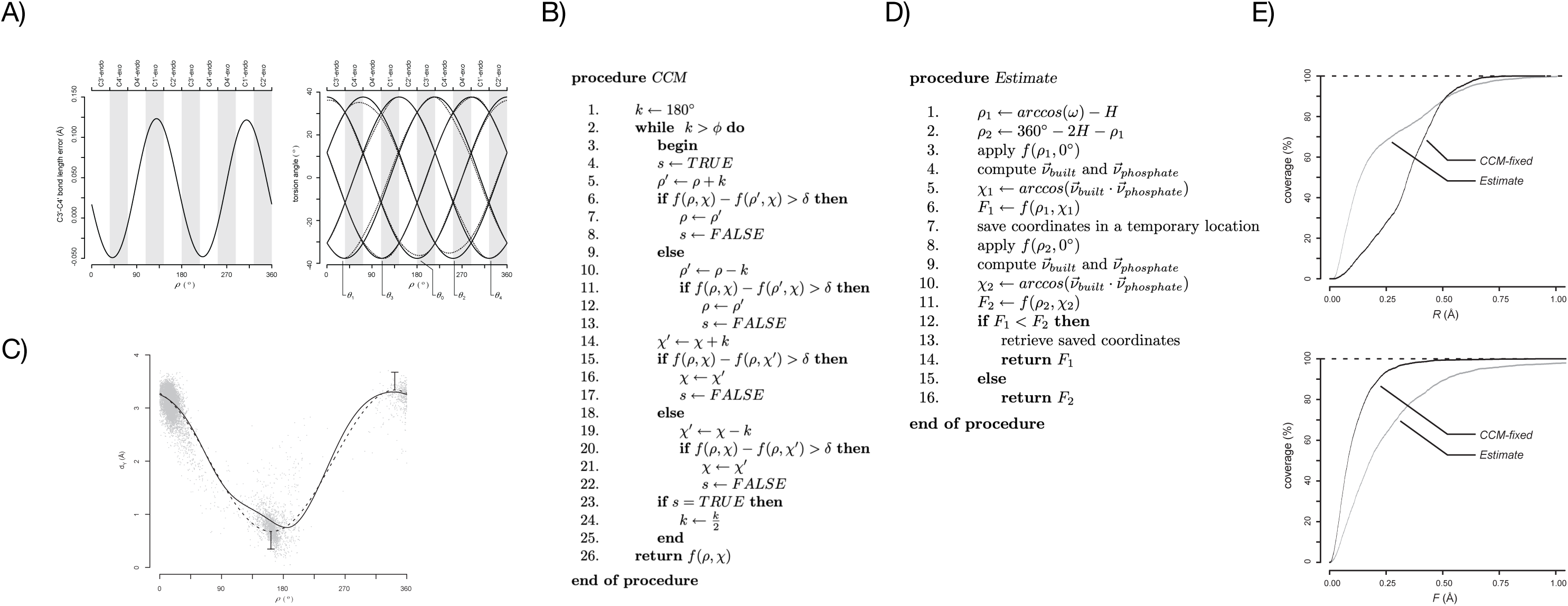
Procedures and precision. **A**) Objective function precision. **Left**) C3’C4’ bond length error against ρ in ribose built by the objective function. **Right**) Error in the furanose torsion angles. The solid lines show the values of the five torsions as computed by Equation 1 across a 360° period for ρ, while the dashed lines shows the values measured on the built ribose. **B**) CCM pseudocode. **C**) Relationship between ρ and O3’. Here *dY* is plotted in solid line against ρ for riboses built across a 360° period for ρ, while χ is kept at 0°. Equation 5 is superimposed in dashed line, with its extrema extended by 0.33 Å (see error bars) to illustrate the effect of accepting arccosine arguments between 1.25 and 1.25. The gray dots show the same plot for the riboses in Σ. **D**) Estimate pseudocode. **E**) Empirical cumulative distribution functions. **Top**) The RMSD between the riboses in Σ’ and those in Σ’C (solid line) and Σ’E (dashed line). **Bottom**) The minimal value of *f*(ρ, χ) after optimization in Σ’C (solid line) and Σ’E (dashed line).

### Optimization

In this section, we introduce two optimization methods that utilize *f*(ρ, χ) as the guiding objective function. By pinpointing the best values for ρ and χ that yield the lowest *f*(ρ, χ), as outlined in Equation 4, we can craft the most suitable ribose conformation that bridges a base with two phosphate groups. The two optimization techniques we employed were: (1) the cyclic coordinate method, and (2) an optimal parameter estimation approach.

#### Cyclic coordinate method

We employed the cyclic coordinate method (20) (CCM), a multidimensional search technique that does not rely on the analytical derivatives of the function. In essence, the CCM algorithm optimizes one parameter at a time while maintaining the others at constant values. This process iteratively cycles through all the parameters, narrowing in on their local minima until the function’s reduction is beneath a specified threshold, Δ. Given that we do not possess an analytical representation for *f*(ρ, χ), we incorporated a line search. This search begins with a predetermined step size in each dimension and diminishes the step whenever no further minimization of the objective function is possible in any of the dimensions. The search halts once the reduction falls below another set threshold, φ. We’ve detailed our CCM algorithm in the pseudocode provided in Figure 2B. The reduction threshold, Δ, is strategically employed to prioritize directions with steeper gradients (either positive or negative increments). Though not detailed in this section, we conducted benchmark tests to determine the ideal values for φ and Δ, which were found to be 0.1° and 0.003 Å, respectively.

Within the CCM, every invocation of *f*(ρ, χ) leads to the complete assembly of a ribose conformation, as previously described. As such, a suitable measure of the algorithm’s complexity would be based on the number of calls made to *f*(ρ, χ), which equates to the number of ribose structures generated. However, predicting the number of iterations CCM requires to pinpoint a minimum is challenging. The functional landscape of the equation, *z* = *f*(ρ, χ), only becomes apparent after the ribose structure is constructed. Furthermore, the minimum that CCM identifies is inherently local, and the method lacks a mechanism to navigate between different local minima.

#### Optimal parameter estimation

In the realm of computer modeling, it is not uncommon to reconstruct the same ribose multiple times. The construction method might, for example, be utilized as a subroutine to complete RNA structures that are only partially assembled. A specific fragment – consisting of a base and two phosphate groups – might be repositioned in various configurations throughout the exploration of the conformational search space, as seen in tools like *MC-Sym*. Consequently, the complexity of an optimization method integrating the ribose construction is compounded within the broader complexity of the entire modeling system. To mitigate its impact on modeling efficiency, we devised an optimal parameter estimation method. This approach necessitates invoking the objective function precisely twice for each iteration.

Our estimation method capitalizes on the observation that the furanose ring is directly covalently bonded to one of the phosphate groups via the C3’-O3’ bond. As a result, the O3’ atom can be constructed as an exocyclic substituent, similar to atoms O2’ and C5’, and is incorporated into the ribose conformation generated by *f*(ρ, χ). Through our investigations, we discerned a relationship between the parameters ρ and χ and the position of O3’ within the ribose. This relationship allows us to predict the values of both parameters based on the provided O3’ position on the 3’ end of the strand.

To facilitate this, we measure the O3’ distance relative to an axis that aligns with the glycosyl bond. Notably, this measurement is solely influenced by ρ and remains unaffected by changes in χ. For our calculations, we selected a coordinate frame where the Y-axis is aligned with the glycosyl bond and the Z-axis is perpendicular to the base plane. Within this framework, the O3’ distance to the glycosyl bond is gauged by its proximity to the Y-axis (refer to dY in Figure 1E).

In Figure 2C, dY is plotted (depicted by a solid line) against varying O3’ exocyclic positions determined by *f*(ρ, χ). This is visualized over a full 360° cycle for ρ, with χ consistently held at 0°. The resulting curve can be closely approximated by a cosine function, as presented in Equation 5. In this equation, *M* signifies the amplitude, *H* represents the phase angle, and *V* denotes the vertical displacement.

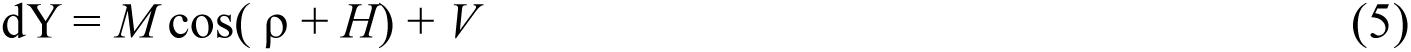

Through our analysis, we established values for the parameters in Equation 5: *M* = 1.33, *H* = 17.42°, and *V* = 2.08 Å. These values optimize the alignment between the observed dY and the cosine function from Equation 5, resulting in a minimal RMSD of 0.115 Å. Additionally, when comparing dY values obtained from riboses in our reference set Σ (as shown by the gray dots in Figure 2C) to those predicted by Equation 5, the RMSD is 0.198 Å. Given the context of molecular modeling, this represents a high level of accuracy. Consequently, we can accurately estimate the value for ρ from dY by rearranging and solving Equation 5.

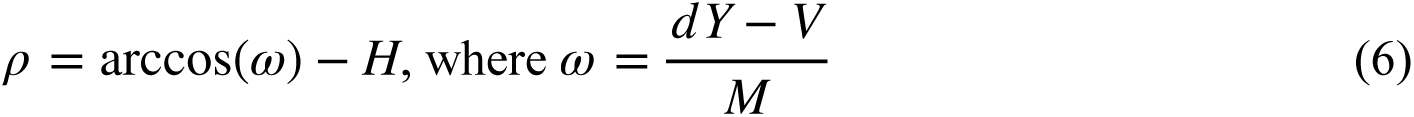

For the successful application of Equation 6, two essential considerations arise.

First, when dealing with the inverse cosine function, one input can yield two distinct angle outcomes, separated by half of a periodic cycle. This duality means that we invariably face a choice between two probable estimates for ρ.

Second, for arccos(ω) to be valid, the constraint -1 ≤ ω ≤ 1 must hold true. When rearranging this for dY, the criteria morph into C – A ≤ dY ≤ C + A. Notably, this condition is not met by 17.5% of the riboses in set Σ. In our bid to bolster its applicability, we adjusted the ω value to either 1 or -1 for values that ranged up to 1.25 or down to -1.25, respectively (as visualized in Figure 2C). This adjustment means that now, a mere 1.6% of riboses in Σ fall outside the stipulated range.

Once we obtain an estimate for ρ, determining χ becomes straightforward. We assemble a ribose, including its exocyclic O3’, using the ρ value estimated from Equation 6 while setting the χ parameter to 0°. On this provisional ribose conformation, we identify two vectors: v_built_ and v_phosphate_. Both vectors are perpendicular to an axis that is aligned with the glycosyl bond. The v_built_ vector originates from the exocyclic O3’ atom of the interim ribose. In contrast, the v_phosphate_ vector is anchored at the corresponding O3’ atom of the 3’ phosphate group (as illustrated in Figure 1F).

The angle formed between the vectors v_built_ and v_phosphate_ reflects the rotation needed around the glycosyl axis to align the intermediate ribose in such a way that the two O3’ atoms — one from the ribose and the other from the 3’ phosphate group — coincide when viewed on a plane perpendicular to the glycosyl axis. In this specific coordinate framework, this rotation corresponds to the value of χ. Given that both v_built_ and v_phosphate_ are unit vectors, we can employ Equation 7 to derive an estimate for χ.

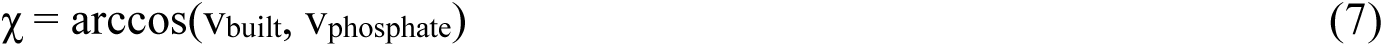

With our methods, we have derived estimates for both ρ and χ. However, it is important to note that our method for estimating ρ provides two mirrored values. To discern the correct ρ value, we compute χ for both potential values. The pair of parameters (ρ and χ) that results in the lowest *f*(ρ, χ) is retained. The comprehensive estimation approach, labeled as “Estimate,” is succinctly presented in the pseudocode depicted in Figure 2D.

Our method consistently requires a fixed four calls to *f*(ρ, χ) for any input to address the ribose construction problem. However, with some minor adjustments, we managed to cut this number down to two. In the procedures at lines 6 and 11, instead of making another call to *f*(ρ, χ) for the determined parameter pair, we apply a rotation transformation to the ribose constructed at lines 3 and 8 to represent the estimated χ. We then directly compute the outcome of Equation 4 to acquire F1 and F2 values. The constant-time complexity of this method stands in stark contrast to that of CCM, where the number of calls to *f*(ρ, χ) is contingent upon the iterations required to find a local minimum.

#### Availability

The two optimization techniques, CCM and Estimate, have been incorporated into the MC-Core bioinformatics development toolkit. *MC-Core* is a C++ object library that we have developed in our laboratory. It is available on the SourceForge open-source code repository, and it is under the MIT License. For individuals with a basic understanding of C++ programming, crafting a small program using the MC-Core library objects to construct ribose conformations becomes a straightforward task. For further guidance, there is a C++ source code provided in the Supporting Information (accessible at no cost). This code allows users to load a PDB-formatted file, recreate the riboses utilizing the Estimate method, and then save the revised coordinates in an output PDB-formatted file. It is essential to note that to compile this code successfully, a proper installation of the *MC-Core* library, version 1.5 or higher, is required.

## Results & Discussion

In this section, we will assess and contrast the performance and precision of the two optimization methods using a benchmark test. We’ll start by introducing the benchmark experiment. Following that, we will delve into an application of the ribose construction method within the computer modeling software *MC-Sym*. We will demonstrate how incorporating the backbone construction post-hoc significantly enhances the accuracy of the generated RNA 3D models. Moreover, this approach substantially reduces the size of the conformational search space being explored.

### Benchmark

Our benchmark involves reconstructing the riboses of the RNA structure found in Σ (see Computational Methods). Initially, we extracted the 17,749 nucleotides from Σ. We then filtered out nucleotides that have an RMSD (Root Mean Square Deviation) of less than 0.25 Å when compared to any other nucleotide. This process yielded a filtered set, denoted as Σ’, containing 3,338 unique nucleotides.

To compare two nucleotide conformations, we employed a specialized distance metric developed in our lab. This metric involves first aligning the atoms in the base, followed by calculating the RMSD between each atom in the backbone (both in the ribose and the phosphate groups) (14).

The filtering process prioritizes the exclusion of nucleotides that have the most neighbors. In this context, two nucleotides are designated as ’neighbors’ if their distance apart is less than the established threshold, which in this case is 0.25 Å. Through this filtering mechanism, we ensure that the benchmark is free from data overrepresentation and its associated biases. One such bias is the RNA ribose pucker’s tendency towards the C3’-endo and C2’-endo conformations (21).

To measure the reduction in pucker overrepresentation within Σ’, we analyzed the distributions of nucleotide conformation types in both Σ and Σ’. A nucleotide conformation type encompasses one of the 10 sugar puckering modes combined with either the syn or anti glycosyl torsions (1). We systematically annotated the nucleotide conformation types using the *MC-Core* library.

Figure S1 presents histograms that compare the nucleotide type distributions in both Σ and Σ’. It is evident from these visual representations that the ribose conformations in Σ’ are more evenly distributed than in Σ. Notably, the dominance of the C3’-endo anti conformation type in Σ is diminished in Σ’, even though it remains relatively prevalent, constituting 56% of the distribution.

To benchmark the ribose construction methods, we removed the ribose atoms from the nucleotides in Σ’, leaving only the bases and phosphate groups to serve as input. The newly constructed riboses were then stored in two sets: Σ’C for the cyclic-coordinate optimization method and Σ’E for the estimation method. The results of this benchmark test are summarized in **Table 1**.

**Table 1:** Benchmark results. The first column, R, shows the expected RMSD between a structure in Σ’ and its rebuilt counterpart in Σ’C and Σ’E. The second and third columns, Δρ, and Δχ, show the expected deviation for ρ and χ, respectively, between their measurement before and after the construction. The fourth column, F, shows the expected, post-optimization, minimal value for *f*(ρ, χ), that is the average bond length error that connects to the phosphate groups. The fifth column, C, shows the expected number of calls to *f*(ρ, χ) needed to rebuild a ribose in Σ’. The proportion of Σ’ that showed better results when rebuilt using *Estimate* is shown in the last row, “*Estimate* gain”. Finally, the last column, N, shows the proportion of Σ’ that was successfully rebuilt.

| | $\overline{R}$ (Å) | $\overline{\Delta\rho}$ (°) | $\overline{\Delta\chi}$ (°) | $\overline{F}$ (Å) | $\overline{C}$ | $N$ (%) |
| --- | --- | --- | --- | --- | --- | --- |
| *CCM* ($\Sigma'C$) | 0.338 | 60.5 | 14.9 | 0.107 | 30 | 100.0 |
| *Estimate* ($\Sigma'E$) | 0.223 | 23.4 | 9.3 | 0.257 | 2 | 98.4 |
| *Estimate* gain (%) | 72.9 | 75.7 | 75.9 | 24.8 | 100.0 |  |

Overall, the Estimate method proved to be more accurate than the CCM in terms of the features R, Δρ, and Δχ (as shown in the first three columns), with superior performance in approximately three out of every four instances. However, in about 75% of cases, the Estimate method produced longer bonds connecting to the phosphate groups. On average, these bonds were around 1.5 times longer than those created by CCM.

Figure 2E showcases the empirical cumulative distribution functions for the RMSD between the original riboses in Σ’ and those in Σ’C and Σ’E, alongside the minimum value of *f*(ρ, χ) post-optimization. As per the RMSD, the Estimate method displayed tighter values, reaffirming the results presented in **Table 1**. Regarding the minimal *f*(ρ, χ), the Estimate method generally settled the optimization issue with marginally higher values. Nonetheless, these remained within acceptable limits, with only 2.2% of the riboses in Σ’E having outliers exceeding 1 Å.

In conclusion, when considering the number of calls to *f*(ρ, χ) required to construct a ribose, the Estimate method significantly outperforms CCM in terms of efficiency without compromising on precision. Specifically, CCM required between 15 to 55 calls, highlighting the efficiency of Estimate.

### Backbone modeling

In this section, we discuss the results achieved by incorporating the ribose construction technique into *MC-Sym*. *MC-Sym* generates RNA 3D structures by systematically assembling nucleotides in 3D space. This positioning is achieved through the application of pre-computed relations for base stacking and base pairing, which are represented as rigid body linear transformations (i.e., translations and rotations).

These relations are derived from high-resolution X-ray crystal structures and have been cataloged in a database in the form of transformation matrices. When modeling a specific RNA, the software maps interactions within that RNA to the appropriate matrices from the database. These matrices then guide the step-by-step placement and transformation of nucleotides.

*MC-Sym* utilizes complete nucleotides, positioning them through the application of these transformation matrices. To maintain the coherency of the backbone structure, the software iteratively tries out multiple nucleotides in different orientations. Only those nucleotides that adhere to the adjacency constraint (specifically, the O3’-P bond length falling within an acceptable range around 1.6 Å) are retained in the model (12).

Figure 1G provides a visual representation of the *MC-Sym* process for creating RNA 3-D models. The example highlights the placement of two adjacent, stacked guanine (G) nucleotides. For this demonstration, we have a set, C, of rigid G nucleotides and another set, T, of transformation matrices corresponding to the stacking relation.

To generate a model, two nucleotides are selected (*a* and *b* from set C) along with a transformation matrix (*t* from set T). The nucleotide ’*a*’ serves as the reference frame for this modeling exercise, and nucleotide ’*b*’ is positioned relative to ’*a*’ using the selected transformation matrix ’*t*’. Given these parameters, there are |C|²|T| potential models that can be generated for this simplistic problem. These models are then refined based on criteria like the adjacency constraint and avoidance of atomic collisions.

In the scenario depicted in Figure 1G-left, the chosen nucleotides are linked by an O3’-P bond with an acceptable length of 1.6 Å when ’*b*’ is positioned relative to ’*a*’ using the transformation ’*t*’. Conversely, Figure 1G-right illustrates a situation where, despite using the same transformation ’*t*’, the choice of different conformational states for nucleotides ’*a*’ and ’*b*’ results in an extended O3’-P bond with an unsuitable length of 7.0 Å.

In practice, it has been recommended to subject *MC-Sym*’s models to energy minimization to rectify backbone inconsistencies (12). One approach to achieve this is by utilizing molecular mechanics, as exemplified in the Amber software package (22). However, this process may inadvertently shift the bases from their designated positions unless properly constrained. An alternative solution, which we introduce here, is the integration of the ribose construction methodology into *MC-Sym*. In the subsequent sections, we will assess and contrast the efficiency and accuracy of models generated by both the original and the updated *MC-Sym* versions. We will not delve into the specific changes made to *MC-Sym* in this context.

Consider the task of constructing two adjacent and stacked G nucleotides once more. Using the terminology defined earlier, we employed a set, C, consisting of 1,100 G nucleotide conformations, and a set, T, containing 1,284 transformation matrices. The original *MC-Sym* is capable of producing approximately 1.55 x 10^9^ different models. With the integration of the ribose construction method, the set C is rendered unnecessary. This is because only the application of the transformation matrices to the rigid body configurations of the bases and phosphate groups is required to produce models. As a result, the number of produced models drops significantly to just 1,284, equivalent to the size of set T. It’s important to note that the additional models generated by the original *MC-Sym* only vary in terms of their backbone structures.

To gauge the quality of the backbone structures, we examined several factors. In the models produced by the original *MC-Sym*, the integrity of the backbone is assessed based on the lengths of the connecting O3’-P bonds, which serve as the adjacency constraint. In contrast, for models generated using the ribose construction method, the quality is measured by averaging the lengths of the implicit bonds that connect both ribose formations to the phosphate group linked to the second nucleotide. **Figure S2** presents histograms illustrating the quality of the backbone structures for all produced models based on the aforementioned measures.

Incorporating the ribose construction method has proven markedly more efficient than the approach of rigid nucleotide sampling used in the original software. When ribose construction is utilized, specifically with the CCM and Estimate methods, 97.9% and 78.7% of the generated models respectively possess a backbone connection within the range of 1 to 2 Å. In stark contrast, only 17.9% of the models produced by the original *MC-Sym* fit this criteria. Alarmingly, in the original models, the backbone connection stretches beyond 5 Å in 26.8% of cases, and in some instances, extends close to 20 Å (not depicted in the data presented).

The histograms from **Figure S2** reveal that the CCM method offers better accuracy compared to the Estimate method. However, this comes at a computational cost: CCM required 76,818 calls to the objective function to produce the 1,284 models. In contrast, the Estimate method necessitated only 5,136 calls, which translates to a time-saving of 1,500% in ribose construction. This significant time efficiency aligns with the benchmark results discussed earlier.

### Ribose stereochemistry

In the ribose construction method, we held all covalent bond lengths and angles constant, characterizing the structure primarily by its torsion angles. However, one essential structural feature of ribose we assumed to be constant was its stereochemistry.

The synthesis of furanose can produce sixteen different stereoisomers. This variation arises because each of the four exocyclic substituents (base, O2’, O3’, and C5’) can project from either side of the furanose ring. The parent molecule of furanose can be any of the four pentoses: ribose, arabinose, xylose, and lyxose. These pentoses differ based on the relative position of their hydroxyl groups across their three chiral carbon atoms. Additionally, each pentose has an enantiomeric configuration labeled as ‘D’ or ‘L’.

As a pentose forms a cyclic structure and becomes a furanose through hemiacetal formation, the type of pentose dictates the orientation of the exocyclic substituent. At the cycle’s junction, rotations at the anomeric carbon classify the resulting furanose stereochemistry into α or β variants. Therefore, the sixteen possible furanose stereoisomers fall into one of four pentose categories, each with two enantiomeric configurations (L or D) and two anomeric types (α or β).

In RNA molecules, riboses are typically of the β-D-ribofuranose type. **Figure S3** visually demonstrates the stereochemistry of this standard ribose variant and contrasts it with three other potential types from the total of sixteen, emphasizing the differences in the relative positions of the exocyclic substituents.

In our ribose construction method, we specifically built the standard β-D-ribofuranose stereoisomer, which is the form typically observed in RNA structures. Yet, within our reference set Σ of X-ray crystallographic structures, a few anomalies were found. Out of 17,749 nucleotides, we identified 27 that did not conform to the standard furanose type: 22 were β-D-xylofuranoses (**Figure S3b**); one was a β-D-arabinofuranose (**Figure S3c**); and, four were α-D-ribofuranoses (**Figure S3d**).

These anomalies were excluded from our benchmark experiment. However, we were curious if these uncommon riboses held functional significance in RNA or if they were merely artifacts from the X-ray crystallographic data capture process.

We employed the CCM method to reconstruct the 27 non-standard riboses into the typical β-D-ribofuranose form. The construction error, represented as F, which corresponds to the optimal outcome of *f*(ρ, χ), is outlined in **Table 2**.

**Table 2:** Errors in rebuilding atypical riboses using the *CCM* method. ^1^Fragment from the human immunodeficiency virus type 1 (HIV-1) Rev response element (RRE). ^2^*Escherichia coli* phage MS2 RNA hairpin-coat-protein complex. ^3^RNA aptamer binded with vitamin B12. ^4^*Haloarcula marismortui* 23S ribosomal subunit. ^5^RNA oligonucleotides binded to RNA polymerase in bacteriophage φ6.

| PDB code | nucleotide | furanose type | $F$ (Å) |
| --- | --- | --- | --- |
| 1CSL<sup>1</sup> | B78 | β-D-arabinofuranoside | 0.417 |
| 1E6T<sup>2</sup> | R2 | β-D-xylofuranoside | 0.141 |
| | S2 | β-D-xylofuranoside | 0.055 |
| 1ET4<sup>3</sup> | A212 | β-D-xylofuranoside | 0.103 |
| | B312 | β-D-xylofuranoside | 0.209 |
| | B323 | β-D-xylofuranoside | 0.015 |
| | C412 | β-D-xylofuranoside | 0.177 |
| | D512 | β-D-xylofuranoside | 0.134 |
| | D523 | β-D-xylofuranoside | 0.080 |
| | E112 | β-D-xylofuranoside | 0.048 |
| 1FFK<sup>4</sup> | '0'282 | β-D-xylofuranoside | 0.063 |
| | '0'600 | β-D-xylofuranoside | 0.030 |
| | '0'894 | β-D-xylofuranoside | 0.065 |
| | '0'904 | β-D-xylofuranoside | 0.045 |
| | '0'1309 | β-D-xylofuranoside | 0.033 |
| | '0'1699 | β-D-xylofuranoside | 0.100 |
| | '0'1835 | β-D-xylofuranoside | 0.050 |
| | '0'1853 | β-D-xylofuranoside | 0.017 |
| | '0'1907 | β-D-xylofuranoside | 0.042 |
| | '0'2083 | β-D-xylofuranoside | 0.051 |
| | '0'2427 | β-D-xylofuranoside | 0.010 |
| | '0'2692 | α-D-ribofuranoside | 1.846 |
| | '0'2714 | β-D-xylofuranoside | 0.046 |
| | '0'2749 | β-D-xylofuranoside | 0.022 |
| 1UVM<sup>5</sup> | D4 | α-D-ribofuranoside | 1.627 |
| | E4 | α-D-ribofuranoside | 1.511 |
| | F4 | α-D-ribofuranoside | 1.464 |

The 22 instances of β-D-xylofuranose were successfully reconstructed into the standard β-D-ribofuranose, with construction errors ranging from 0.010 Å to 0.209 Å. **Figure S4** displays a successful transformation of the β-D-xylofuranose ribose found in nucleotide ’0’1853 of the PDB entry 1FFK.

The singular β-D-arabinofuranose had a slightly elevated error of 0.417 Å. However, the four α-D-ribofuranoses posed a more significant challenge, with errors ranging from 1.464 Å to 1.846 Å. As demonstrated in **Figure S5**, the attempt to transform an α-D-ribofuranose resulted in a distorted ribose structure. As a result, our method could not adequately transform these α-D-ribofuranoses into the standard β-D-ribofuranose structure.

## Conclusions

We designed and executed methods for constructing RNA riboses that link a base to its two neighboring phosphate groups. These techniques are both highly effective and beneficial in the realm of RNA 3D modeling.

We introduced two distinct optimization techniques to address the ribose construction challenge: a linear search, termed CCM, and a fixed-time estimation strategy, named Estimate. Both methods consistently produced ribose conformations that aligned closely with those found in a reference set of X-ray crystallographic structures. The incorporation of the ribose construction technique significantly enhances the efficiency of the RNA 3D modeling software. This enhancement arises because the RNA building process is divided into two separate tasks: Positioning the bases and phosphate groups and constructing the ribose conformations.

The first task is now streamlined, as only the bases and phosphate groups need spatial positioning as fixed entities. This simplification allows for a more focused exploration of RNA’s conformational search space. The second task is efficiently tackled using numerical methods.

The benchmarking results indicated that while CCM constructed riboses with fewer errors compared to the Estimate method, it did so at a significantly higher computational expense. Specifically, the CCM approach required a varying and unpredictable number of calls to the function *f*(ρ, χ), whereas the Estimate method consistently used only two calls. Additionally, the original structural characteristics of the riboses in the benchmark experiment were better preserved when using the Estimate approach.

Comparing the accuracy of the two implementations, CCM and Estimate, presents a challenge. Determining whether to prioritize the preservation of structural features or minimizing construction errors is a complex decision. However, in the context of RNA 3D structure modeling, the advantages of using Estimate become more evident due to its constant-time computational efficiency. Specifically, when applied in *MC-Sym* to model two consecutive G nucleotides, Estimate demonstrated a significant speed advantage. It achieved a performance improvement of 1500%, requiring only 5,136 calls to *f*(ρ, χ) to create 1,284 distinct structures, compared to CCM’s 76,818 calls to *f*(ρ, χ).

## Acknowledgment

This work was supported by a grant from the Canadian Institutes of Health Research (CIHR) (MT-14604) to FM. PT had a scholarship from the Fonds de Recherche du Québec sur la Nature et les Technologies.

