## Supplementary Table and Figures for "Building RNA Backbone in Constant Time by Numerical Approximation"

### (Supplementary Materials)

**Philippe Thibault and François Major**

Institute for Research in Immunology and Cancer
Department of Computer Science and Operations Research
Université de Montréal
PO Box 6128, Downtown station
Montréal, QC H3C 3J7
CANADA

### Supplementary Table

**Table**: PDB codes of high-resolution X-ray crystallographic structures that define database Σ.

| | | | | | | | | | | |
|---|---|---|---|---|---|---|---|---|---|---|
| 157D | 165D | 1A34 | 1A9N | 1C0A | 1C9S | 1CSL | 1CX0 | 1D4R | 1D96 | 1DFU |
| 1DI2 | 1DNO | 1DNT | 1DQF | 1DQH | 1DRZ | 1DUL | 1DUQ | 1E6T | 1E7X | 1EC6 |
| 1EFO | 1EHZ | 1ET4 | 1EVP | 1EVV | 1F27 | 1F7U | 1FEU | 1FFK | 1FFY | 1FIX |
| 1FUF | 1FXL | 1G2E | 1G2J | 1G4Q | 1G59 | 1GTF | 1H2C | 1HQ1 | 1HR2 | 1I2X |
| 1I2Y | 1I9X | 1ICG | 1ID9 | 1IDW | 1IHA | 1IK5 | 1J1U | 1J6S | 1J8G | 1J9H |
| 1JB8 | 1JBR | 1JBS | 1JID | 1JJ2 | 1JO2 | 1JZV | 1K8W | 1KD3 | 1KD4 | 1KD5 |
| 1KFO | 1L2X | 1L3Z | 1LNG | 1LNT | 1M5K | 1M5O | 1M5V | 1M8W | 1M8X | 1MDG |
| 1MSW | 1MSY | 1MWL | 1N77 | 1N78 | 1N7A | 1N7B | 1NLC | 1NUJ | 1NUV | 1O9M |
| 1OFX | 1OSU | 1P79 | 1PJG | 1PJO | 1Q93 | 1Q96 | 1Q9A | 1QBP | 1QC0 | 1QLN |
| 1QTQ | 1QU2 | 1R3E | 1R9F | 1RC7 | 1RNA | 1RXB | 1S72 | 1T0D | 1T0E | 1TFW |
| 1U0B | 1U8D | 1URN | 1UTD | 1UTF | 1UTV | 1UVI | 1UVJ | 1UVM | 1W55 | 1WMQ |
| 1WSU | 1Y26 | 1Y27 | 1Y99 | 1YHQ | 246D | 248D | 255D | 259D | 280D | 283D |
| 2A8V | 2BGG | 2BH2 | 315D | 332D | 353D | 354D | 377D | 397D | 398D | 402D |
| 406D | 413D | 418D | 419D | 420D | 421D | 430D | 433D | 434D | 435D | 437D |
| 439D | 462D | 464D | 466D | 472D | 479D | 480D | 483D | 485D | | |

### Supplementary Figures

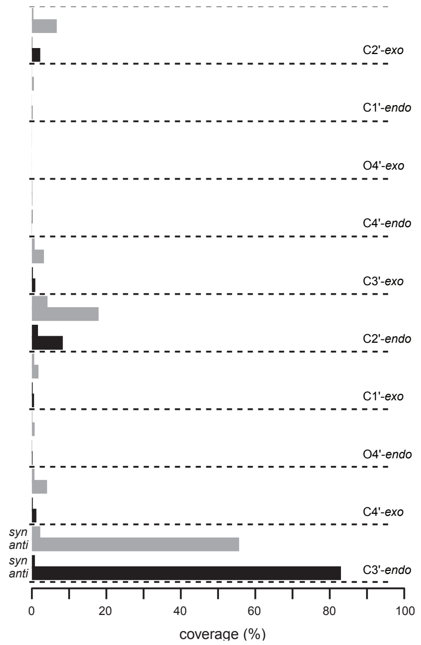


**Figure S1**. **Nucleotide conformational type distributions**. The nucleotide conformations in Σ are shown in black; in Σ' in gray. For each of the 10 puckering modes (delimited by dashed lines), two bars for each set are shown, separating the two glycosyl torsions: *syn* and *anti*.

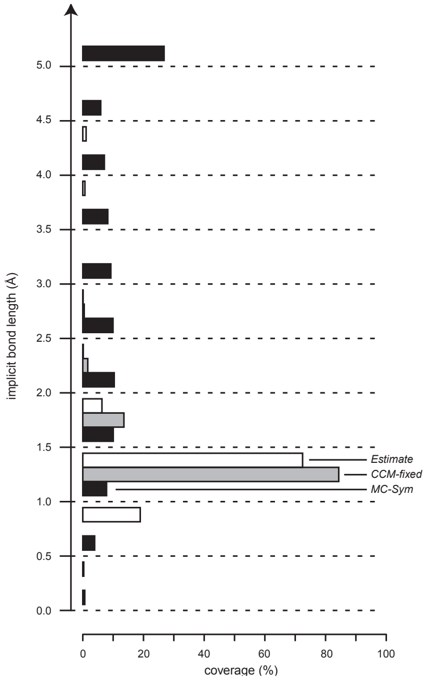


**Figure S2**. **Backbone quality**. The histograms show the backbone quality of two stacked G nucleotides using the original *MC-Sym* (black) and that embedded the ribose construction, using *CCM* (gray) and *Estimate* (white). The lengths of the implicit covalent bonds inherent to each construction method quantify the backbone quality.

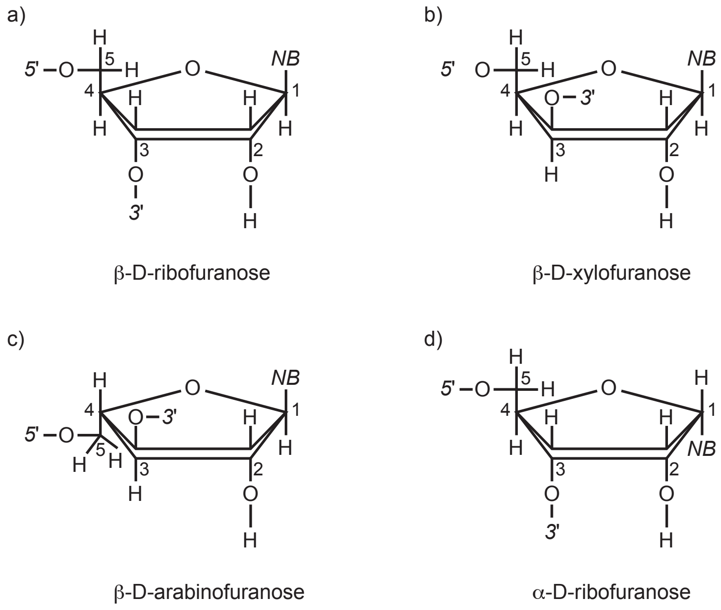


**Figure S3**. **Four furanose types**. The schematized carbons are numbered from 1 to 5. *NB* connects to the base, 5’ to the next phosphate group toward the 5’-end, and 3’ to the next phosphate group toward the 3’-end.

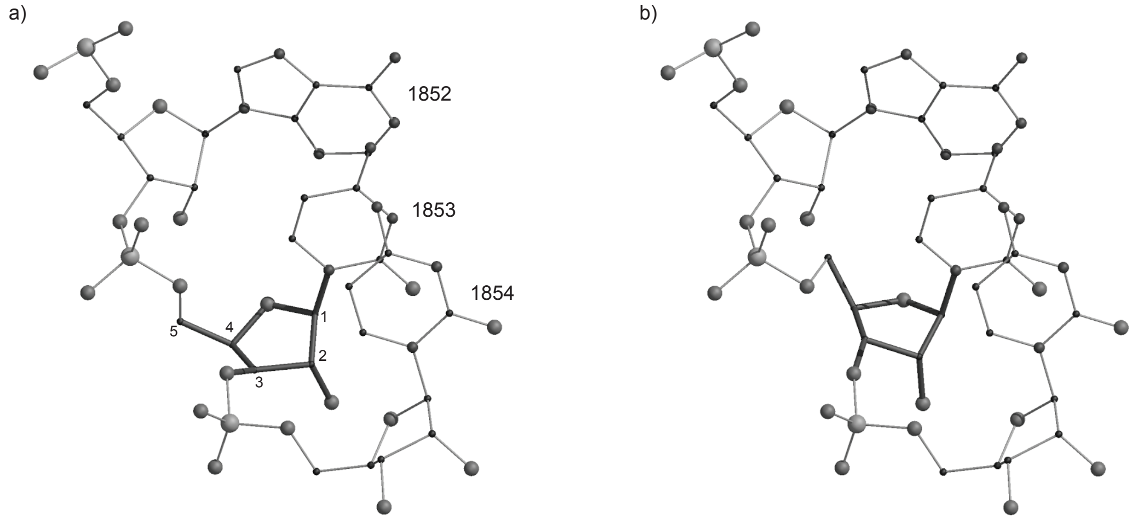


**Figure S4**. **Fragment including a β-D-xylofuranoside**. The fragment is from the 23S ribosomal subunit: 1852-ACC-1854 in chain 0 of PDB entry 1FFK. a) Original fragment. The βDxylofuranoside of nucleotide 1853 is emboldened and its carbon atoms numbered. b) Reconstruction of the ribose in nucleotide 1853 as a β-D-ribofuranoside using *CCM*.

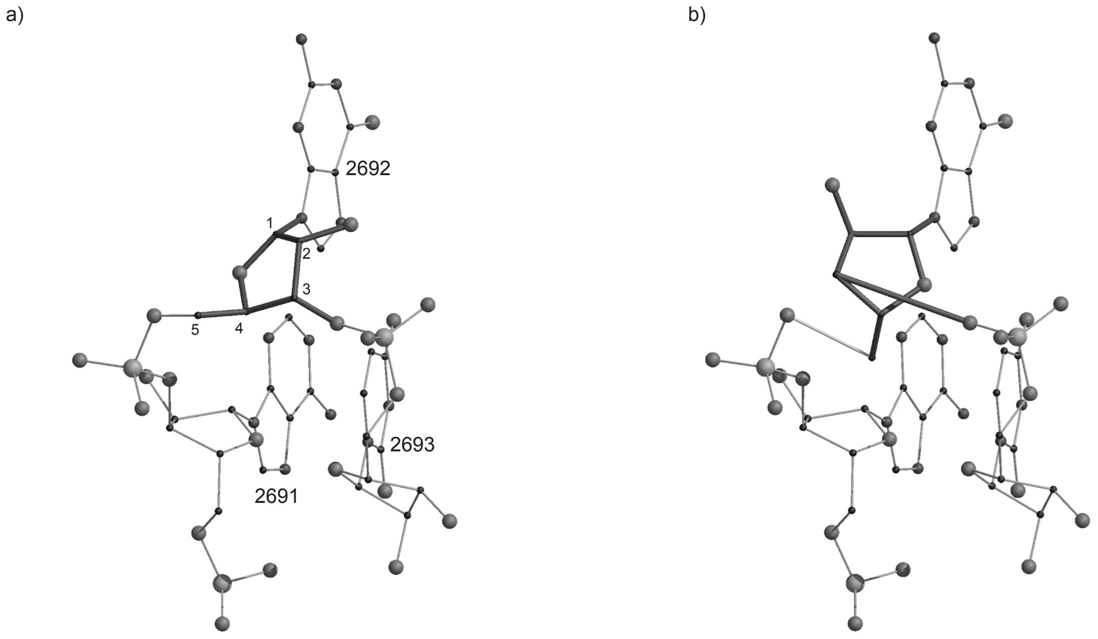


**Figure S5**. **Fragment including an α-D-ribofuranoside**. The fragment is from the 23S ribosomal subunit: 2691-AGU-2693 in chain 0 of PDB entry 1FFK. a) Original fragment. The αDribofuranoside of nucleotide 2692 is emboldened and its carbon atoms numbered. b) Failed reconstruction of the ribose in nucleotide 2692 as a β-D-ribofuranoside using *CCM*.
